# MinION barcodes: biodiversity discovery and identification by everyone, for everyone

**DOI:** 10.1101/2021.03.09.434692

**Authors:** Amrita Srivathsan, Leshon Lee, Kazutaka Katoh, Emily Hartop, Sujatha Narayanan Kutty, Johnathan Wong, Darren Yeo, Rudolf Meier

**Affiliations:** Department of Biological Sciences, National University of Singapore, Singapore; Research Institute for Microbial Diseases, Osaka University, Japan; Artificial Intelligence Research Center, AIST, Tokyo, Japan; Zoology Department, Stockholms Universitet, Stockholm, Sweden; Station Linné, Öland, Sweden; Tropical Marine Science Institute, National University of Singapore, Singapore; Museum für Naturkunde, Leibniz Institute for Evolution and Biodiversity Science, Center for Integrative Biodiversity Discovery, Berlin, Germany

## Abstract

**Background:** DNA barcodes are a useful tool for discovering, understanding, and monitoring biodiversity which are critical tasks at a time of rapid biodiversity loss. However, widespread adoption of barcodes requires cost-effective and simple barcoding methods. We here present a workflow that satisfies these conditions. It was developed via “innovation through subtraction” and thus requires minimal lab equipment, can be learned within days, reduces the barcode sequencing cost to <10 cents, and allows fast turnaround from specimen to sequence by using the portable, real-time sequencer MinION.

**Results:** We describe cost-effective and rapid procedures for barcoding individual specimens with MinION sequencing. We illustrate how tagged amplicons can be obtained and sequenced with the portable, real-time MinION sequencer in many settings (field stations, biodiversity labs, citizen science labs, schools). We also provide amplicon coverage recommendations that are based on several runs of the latest generation of MinION flow cells (“R10.3”) which suggest that each run can generate barcodes for >10,000 specimens. Next, we present a novel software, ONTbarcoder, which overcomes the bioinformatics challenges posed by MinION reads. The software is compatible with Windows 10, Macintosh, and Linux, has a graphical user interface (GUI), and can generate thousands of barcodes on a standard laptop within hours based on only two input files (FASTQ, demultiplexing file). We document that MinION barcodes are virtually identical to Sanger and Illumina barcodes for the same specimens (>99.99%) and provide evidence that MinION flow cells and reads have improved rapidly since 2018.

**Conclusions:** We propose that barcoding with MinION is the way forward for government agencies, universities, museums, and schools because it combines low consumable and capital cost with scalability. Small projects can use the flow cell dongle (“Flongle”) while large projects can rely on MinION flow cells that can be stopped and re-used after collecting sufficient data for a given project.

## 1. Background

DNA sequences have been used for identification and taxonomic purposes for decades [1–3], but for most of this time they have been akin to mobile phones in the 1990s: of limited value due to sparse signal coverage and high cost. Obtaining barcodes was too complicated and expensive despite the development of effective DNA extraction protocols [4], fast Sanger sequencing protocol [5], and the establishment of the Canadian Centre for DNA Barcoding (CCDB), which is the primary sequencing facility for the International Barcode of Life Consortium (iBOL: https://ibol.org/resources/sequencing-facility/). After 15 years and the investment of >200 million USD in facilities and barcoding, ca. 8 million animal barcodes are available for searches in the database “BOLD Systems” of which ca. 6 million sequences can be downloaded as “public data” (April 2021: [6]). Combined with barcodes from NCBI, they are now a valuable resource for the global biodiversity community.

However, the cost of barcodes has remained high (http://ccdb.ca/pricing/) and the most widely used approach to large-scale barcoding still relies on sending specimens from all over the world to the CCDB in Guelph (Canada). For example, >85% of all arthropod barcodes in BOLD Systems were generated by the CCDB with more than >60% of the voucher specimens remaining in the centre (https://www.boldsystems.org/index.php/Taxbrowser_Taxonpage?taxid=20). Unfortunately, this model for barcoding interferes with real-time biodiversity monitoring and specimen accessibility. We therefore here argue that barcoding has to be decentralized. We show that this can be achieved through “innovation through subtraction,” which yields simplified and cost-effective solutions for generating barcode amplicons in molecular laboratories with very basic equipment. Combined with the use of MinION sequencers, these simplifications allow for generating barcodes almost anywhere by biologists and citizen scientists alike.

A decentralized model for monitoring the world’s biodiversity is necessary given the scale, urgency, and importance of the task at hand. For example, even if there were only 10 million species of metazoan animals on the planet [7] and a new species was discovered with every 50^th^ specimen that is processed, species discovery with barcodes will require the sequencing of 500 million specimens [8]. Yet, species discovery is only a small part of the biodiversity challenge in the 21^st^ century. Biodiversity loss is now considered by the World Economic Forum as one of the top three global risks based on likelihood and impact for the next 10 years [9] and Swiss Re estimates that 20% of all countries face ecosystem collapse as biodiversity declines [10]. This implies that biodiversity discovery and monitoring will require in the future completely different scales than in the past. The old approaches thus need rethinking because all countries will need real-time distributional and abundance information in order to develop effective conservation strategies and policies. In addition, they will need information on how species interact with each other and the environment [11].

Barcodes were initially intended as an identification tool for biologists [1]. Thus, most projects focused on taxa with a large following in biology (e.g., birds, fish, butterflies) [12] Yet, despite targeting taxa with well-understood diversity, the projects struggled with barcoding >75% of the described species in these groups [12]. When the pilot barcoding projects ran out of tissues from identified specimens, they started targeting unidentified specimens; i.e., DNA barcoding morphed into a technique that was used for biodiversity discovery (“dark taxa”: [12, 13]). This shift towards biodiversity discovery was gradual and incomplete because the projects used a “hybrid approach” that started with sorting specimens to “morphospecies” before barcoding one or a few specimens for each morphospecies/sample (e.g., [14–20]). However, this approach is not ideal for biodiversity discovery and monitoring because morphospecies sorting is labour-intensive and of unpredictable quality because it is heavily dependent on the taxonomic expertise of the sorters [21, 22]. This is why it is preferable to reverse the workflow [23] by barcoding all specimens first and assessing congruence with morphology afterwards [24, 25]. This approach has the additional benefit that it yields quantitative data and corroborated species-level units. However, it requires efficient and low-cost barcoding methods that are also suitable for biodiverse countries with limited science funding.

Fortunately, such cost-effective barcoding methods are now becoming available. This is partially due to the replacement of Sanger sequencing with second- and third-generation sequencing technologies that have lower sequencing costs [23, 26–33]. These changes mean that the widespread application of the reverse workflow is now feasible for tackling the species-level diversity of those metazoan clades that are so specimen- and species-rich that they have been neglected in the past [33, 34]. Many of these clades have high spatial species turnover, requiring many localities in each country to be sampled and large numbers of specimens to be processed [32]. Such intensive processing is best achieved close to the collecting locality to avoid the delays and costs of shipping samples across continents. This is now feasible because biodiversity discovery can be readily pursued in decentralized facilities at varied scales. Indeed, accelerated biodiversity discovery is a rare example of a big science initiative that allows for meaningful engagement of students and citizen scientists, which can enhance significantly biodiversity education and appreciation [35–38]. This is especially so when stakeholders not only barcode, but also image specimens, determine species abundances, and map distributions of newly discovered species. All of which can be based on specimens collected in the citizens’ own backyard.

But can such decentralized biodiversity discovery really be effective? Within the last five years, students and interns in the laboratory of the corresponding author at the National University of Singapore barcoded >330,000 specimens. After analyzing the first >140,000 barcoded specimens for selected taxa representing different ecological guilds, the alpha and beta diversity of Singapore’s arthropod fauna was analyzed based on ∼8,000 putative species which revealed that some habitats were unexpectedly species-rich and harboured unique faunas (e.g., mangroves, freshwater swamp: [32, 39]). Barcodes even helped with the conservation of charismatic taxa when they were used to identify the larval habitats for more than half of Singapore’s damsel- and dragonfly species [40] and facilitated species interaction research and biodiversity surveys based on eDNA [41, 42]. Biodiversity appreciation by the public was fostered by featuring newly discovered species and their species interactions on “Biodiversity of Singapore” (BOS >15,000 species: https://singapore.biodiversity.online/), dozens of new species have been described, and the descriptions of another 150 species are being finalized [43–51].

Barcoding metazoan specimens require the successful completion of three steps: (1) obtaining DNA template, (2) amplifying *COI* via PCR, and (3) sequencing the *COI* amplicon. Many biologists learn these techniques in university for a range of different genes – from those that are easy to amplify (short fragments of ribosomal and mitochondrial genes with well-established primers) to those are difficult (long, single-copy nuclear genes with few known primers). Fortunately, amplification of short mitochondrial markers like *COI* does not require the same level of care as nuclear markers. Learning how to barcode efficiently is hence an exercise of unlearning unnecessary procedures. Note that this unlearning comes with cost savings which are particularly vital for boosting biodiversity research where it is most needed: in biodiverse countries with limited science funding. Our manuscript therefore has a more comprehensive Methods section than most publications, because we do not only describe which methods we used for our experiments, but also why certain alternative methods were avoided. In addition, we provide videos that illustrate the techniques (https://www.youtube.com/channel/UC1WowokomhQJRc71FmsUAcg).

Simplified laboratory techniques for obtaining barcode amplicons are important, but they need to be complemented with efficient and cost-effective sequencing techniques. Therefore, we here also test the capabilities of the latest flow cells used in Oxford Nanopore Technologies (ONT) instruments. They have the advantage of being inexpensive and yielding data quickly by passing single-stranded DNA through a nanopore and using the current fluctuations for reconstructing the DNA sequence [52]. ONT’s MinION is especially suitable for decentralized barcoding because it is small and inexpensive. However, its use for barcoding has been unpopular because of low sequence read accuracy (85% - 95%: [52, 53]). This meant that the data had to be analyzed with complex bioinformatics pipelines that are not suitable for widespread use. Recently, three significant changes occurred. Firstly, ONT released a low-cost and capacity flow cell (Flongle) that only has 126 pores (126 channels) instead of the customary 2048 (512 channels) of a full MinION flow cell. We here test whether Flongle is a promising tool for small barcoding projects that need quick turnaround times for a few hundred specimens.

Secondly, ONT released a new flow cell chemistry for full flow cells (“R10.3”) where the nanopores have a dual instead of a single reader-head. Dual reading has altered the read error profile by giving better resolution to homopolymers and improved consensus accuracy [54, 55]. Lastly, ONT released high accuracy (HAC) basecalling (https://nanoporetech.com/about-us/news/new-research-algorithms-yield-accuracy-gains-nanopore-sequencing). All three innovations are here tested using six different amplicon pools of very different sizes (191 - 9,932 specimens). Overall, we find that the innovations were very effective at improving read quality and quantity. This meant that we could develop and release a new bioinformatics software, “ONTbarcoder,” which is fast and user-friendly in that it has a GUI, is compatible with all major operating systems, and does not require the installation of third-party software.

## Results

### Performance of new flow cells and high-accuracy basecalling

We used six pools of amplicons to test the new ONT products. The pools contained amplicons for 191 - 9,932 specimens and were run for 15-49 hours (Table 1). The amplicons were tagged with a 13-bp index to facilitate demultiplexing reads to specimen specific bins. Basecalling the fast5 files using Guppy in MinIT under the high accuracy (HAC) model was still very slow and took 12 days for the largest dataset (*Palaearctic Phoridae (658 bp)* (Table 1). However, it yielded good quality reads that could be demultiplexed at high rates for the four R10.3 MinION datasets (= 30-49%). The only exception was the *Palaearctic Phoridae (313 bp)* dataset (15.5%). Flongle datasets showed overall also lower demultiplexing rates (17-21%).

**Table 1.**
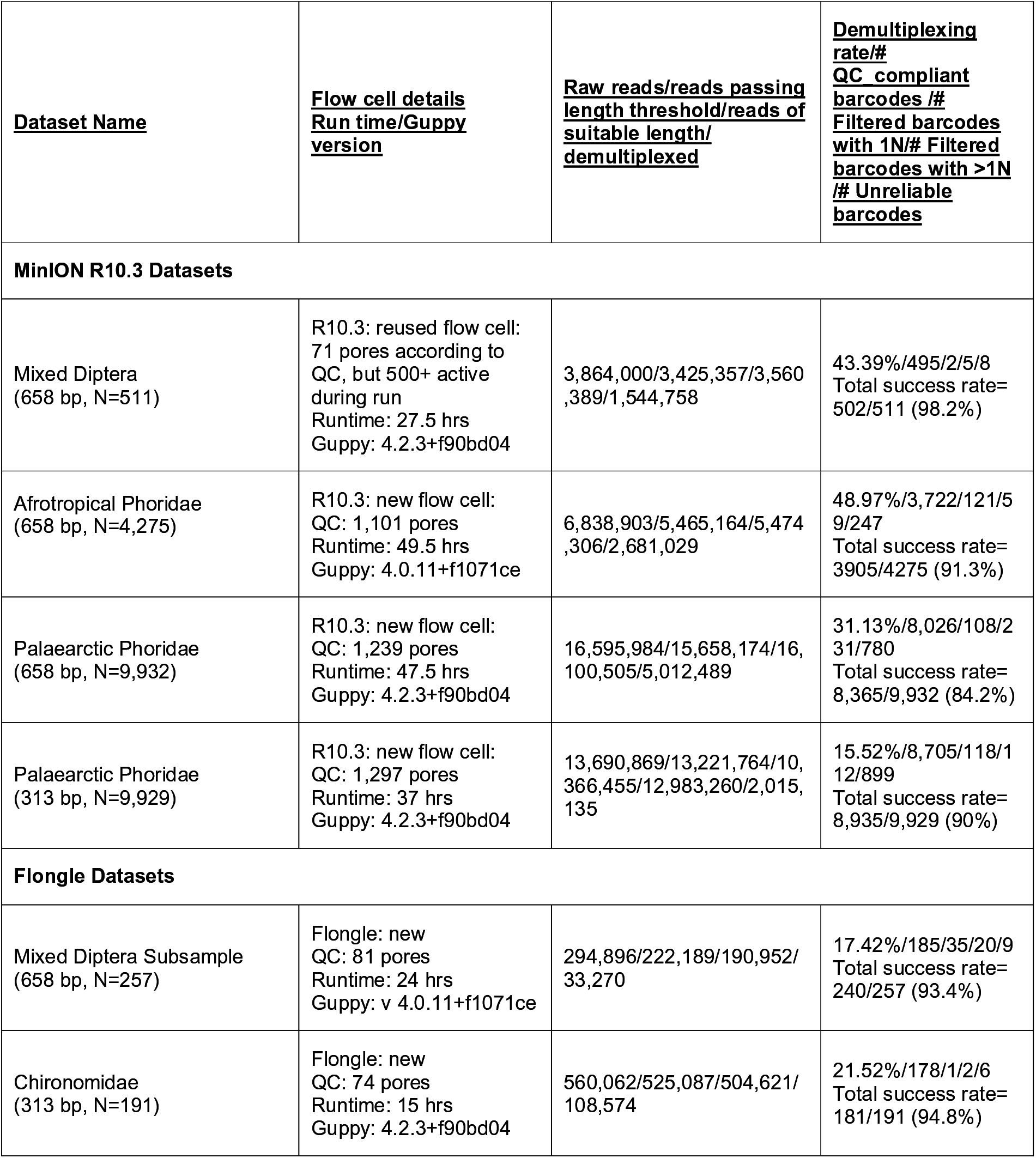
Datasets generated in this study and the results of barcoding using ONTbarcoder at 200X coverage (Consensus by Length) and 100X coverage (Consensus by Similarity).

We then investigated barcode accuracy (Figure 1) by directly aligning the MinION barcodes with Sanger and Illumina barcodes for the same specimens. We find that MinION barcodes are virtually identical (>99.99% identity, Table 2). We furthermore established that the number of ambiguous bases (“N”) is very low for barcodes obtained with R10.3 (<0.01%). Indeed, more than 90% of all barcodes are entirely free of ambiguous bases. In comparison, Flongle barcodes have a slightly higher proportion of ambiguous bases (<0.06%). They are concentrated in ∼20% of all sequences so that 80% of all barcodes again lack Ns. This means that MinION barcodes well exceed the Consortium for the Barcode of Life (CBOL) criteria for “barcode” designation with regard to length, accuracy, and ambiguity (Ratnasingham and Hebert 2013).

**Figure 1.**
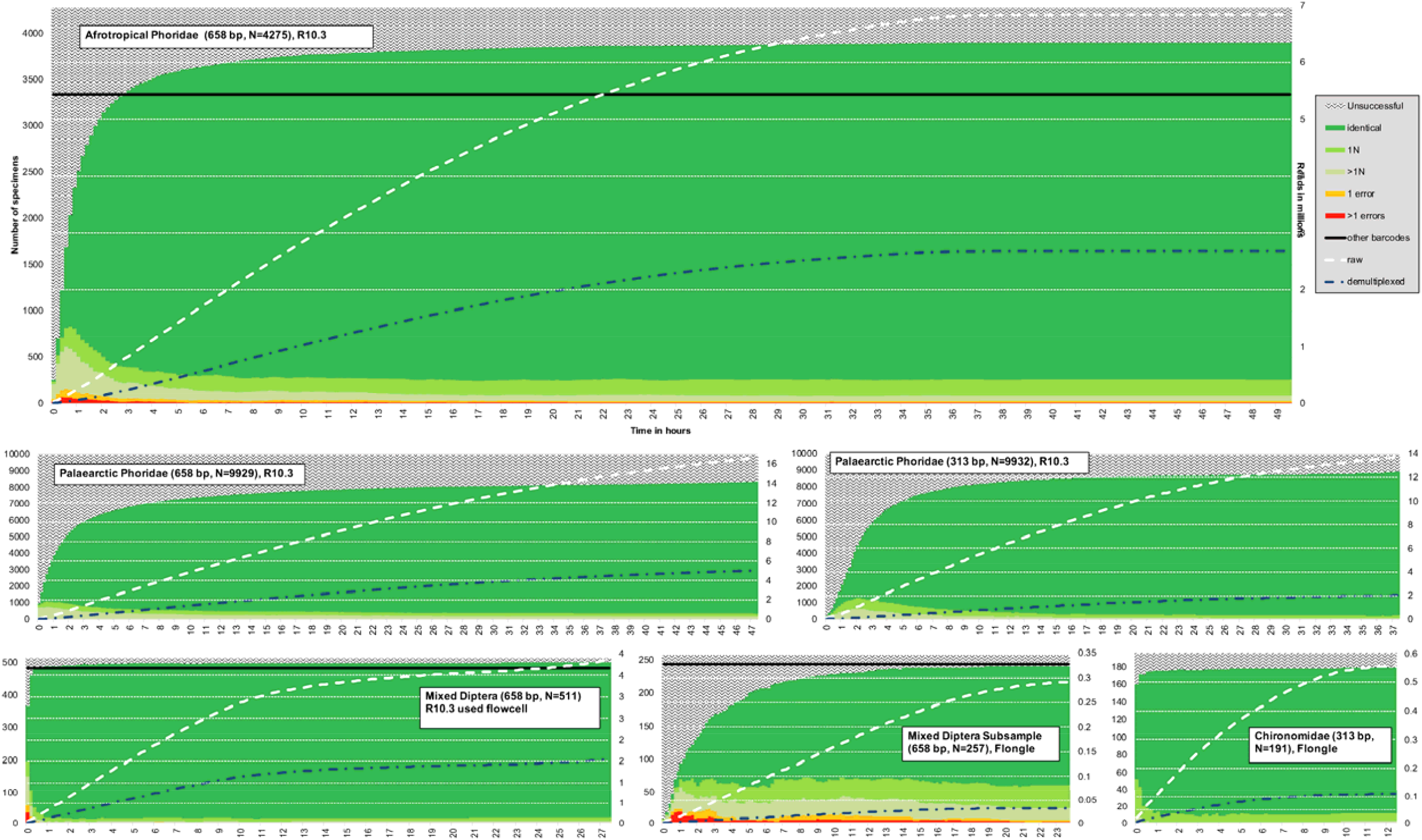
Rapid recovery of accurate MinION barcodes over time (in hours, x-axis) (filtered barcodes: dark green = barcodes passing all 4 QC criteria, light green = one ambiguous base; lighter green = more than 1N, no barcode = white with pattern, 1 mismatch = orange, >1 mismatch = red). The solid black line represents the number of barcodes available for comparison. White dotted line represents the amount of raw reads collected over time, blue represents number of demultiplexed reads over time (plotted against Z-axis)

**Table 2.** Quality assessment of barcodes generated by ONTbarcoder at 200X read coverage (Consensus by Length) and 100X coverage (Consensus by Similarity). The accuracy of MinION barcodes is compared with the barcodes obtained for the same specimens using Illumina/Sanger sequencing. Errors are defined as sum of substitution or indel errors. Denominators are the total number of nucleotides assessed.

| Dataset | No. of comparison barcodes | No. of barcodes with errors/No. of errors/% identity | # of Ns/%Ns |
| --- | --- | --- | --- |
| R10.3: Mixed Diptera: Sanger barcodes available | 476 | 2/10/99.997% | 19 (0.006%) |
| R10.3: Afrotropical Phoridae: Illumina barcodes available* | 3316 | 23/48/99.995% | 284 (0.011%) |
| Flongle-Mixed Diptera Subsample: Sanger barcodes available | 231 | 5/8/99.994% | 91 (0.058%) |

\*5 barcodes with very high distances from reference were excluded for R10.3: Afrotropical Phoridae dataset as they likely represent lab contamination (see Srivathsan, Hartop et al. (2019)

Rarefaction at different read coverage levels reveals that 80-90% of high-quality barcodes are obtained within a few hours of sequencing. In addition, the number of barcodes generated by MinION exceeded or was comparable to what could be obtained with Sanger or Illumina (Figure 1). We also determined the coverage needed for obtaining reliable barcodes. For this purpose, we plotted the number of barcodes obtained against coverage (Figure 2). This revealed that the vast majority of specimens yield high-quality barcodes at coverages between 25x and 50x when R10.3 reads are used. Increasing coverage beyond 50x only leads to modest improvements in quality and few additional barcodes. The coverage needed for obtaining Flongle barcodes was somewhat higher, but the main difference between the R9.4 technology of the Flongle flow cell and R10.3 of the MinION flow cell was that more barcodes retained ambiguous bases even at high coverage for R9.4 data. The differences in read quality between R9.4 and R10.3 became even more obvious when the read bins for the “Mixed Diptera Subsample” were analyzed based an identical numbers of R10.3 and R9.4 reads. The barcodes based on Flongle and R10.3 data are compatible, but the R10.3 barcodes are ambiguity-free while some of the corresponding Flongle barcodes retain 1-2 ambiguous bases.

**Figure 2.**
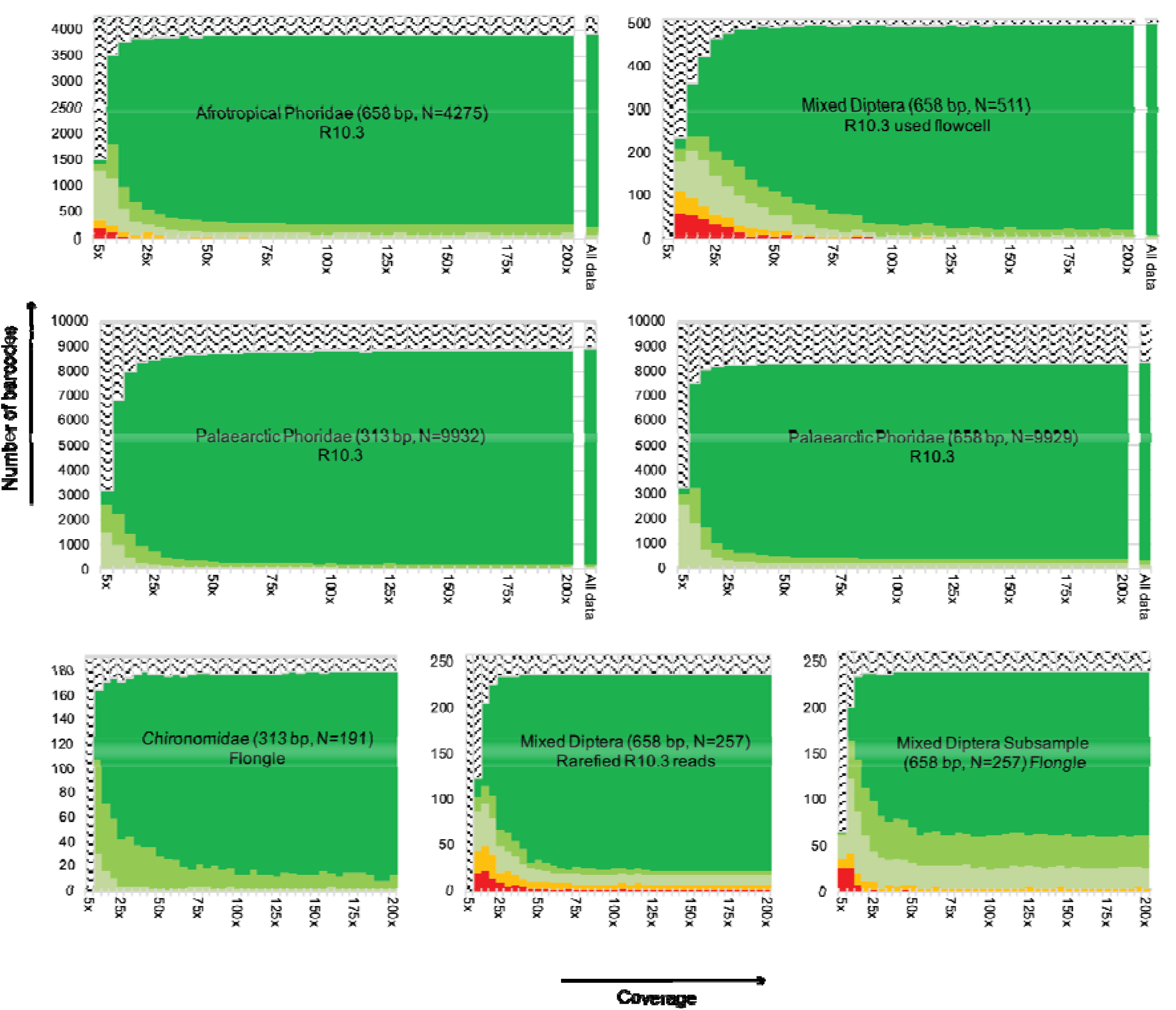
Relationship between barcode quality and coverage. Subsetting the data to 5-200X coverage shows that there are very minor gains to barcode quality after 25-50X coverage. (filtered barcodes: dark green = barcodes passing all 4 QC criteria, light green = one ambiguous base; lighter green = more than 1N, no barcode = white with pattern, 1 mismatch = orange, >1 mismatch = red).

Overall, these results imply that 100x raw read coverage is sufficient for obtaining barcodes with either R10.3 or R9.4 flow cells. Given that most MinION flow cells yield >10 million reads of an appropriate length, one could, in principle, obtain 100,000 barcodes in one flow cell. However, this would require that all amplicons are represented by similar numbers of copies and that all reads could be correctly demultiplexed. In reality, only 30-50% of the reads can be demultiplexed and the number of reads per amplicon fluctuates widely (Figure 3). Very-low coverage bins tend to yield no barcodes or barcodes of lower quality (errors or Ns). These low-coverage barcodes can be improved by collecting more data, but this comes at a high cost and increased risk of a small number of contaminant reads yielding barcodes. For example, we observed that some “negative” PCR controls yielded low-quality barcodes for 4 of 106 negatives in the Palaearctic Phoridae (313 bp) and 1 of 105 negatives in the Palaearctic Phoridae (658 bp) datasets.

**Figure 3.**
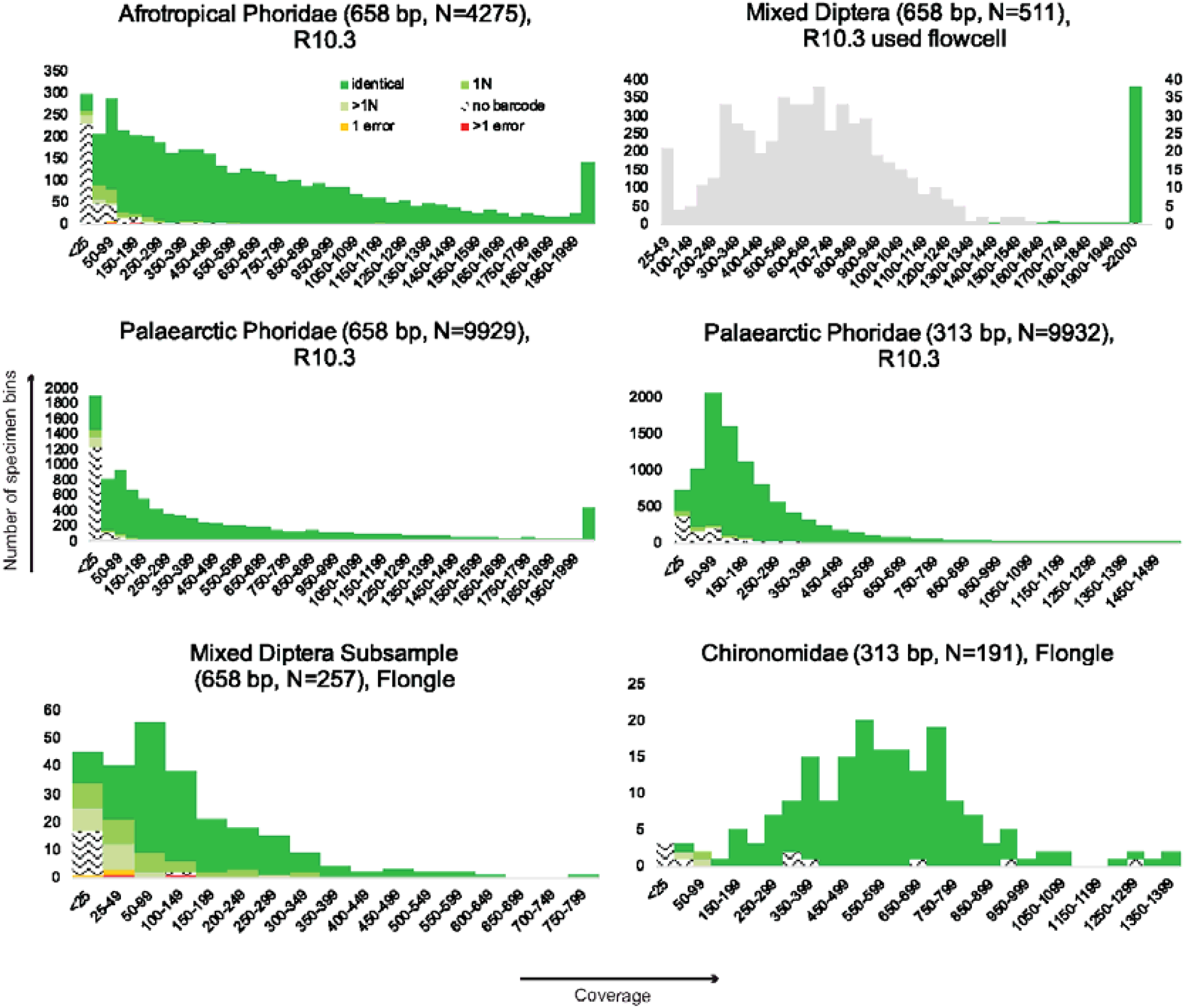
Read bin size distribution for six amplicon pools (color-coding as in Figs 1-2). Due to the very generous coverage for the “Mixed Diptera” dataset, we also use grey to show the bin size distribution after dividing the bin read totals by 5.

To facilitate the planning of barcode projects, we illustrate the trade-offs between barcode yield, sequencing time, and amount of raw data needed for six amplicon pools (Figure 4: 191-9,932 specimens). These standard curves can be used to roughly estimate the number of raw reads needed to achieve a specific goal for a barcoding project of a specific size (e.g., obtaining 80% of all barcodes for a project with 1000 amplicons). Note that the number of raw reads is displayed in real-time in MinKNOW so that the run can be terminated when the target number of reads has been reached. The number of recoverable barcodes in Figure 4 was set to the number of all error-free, filtered barcodes obtained in an analysis of all data. We would argue that this is a realistic estimate of recoverable barcodes given the saturation plots in Figure 1 that suggest that most barcodes with significant amounts of data have been called at 200x coverage. Note, however, that Figure 4 can only provide very rough guidance on how many reads are needed because, for example, the demultiplexing rates differ between flow cells and different amplicon pools have very different read abundance distributions (see Figure 3).

**Figure 4.**
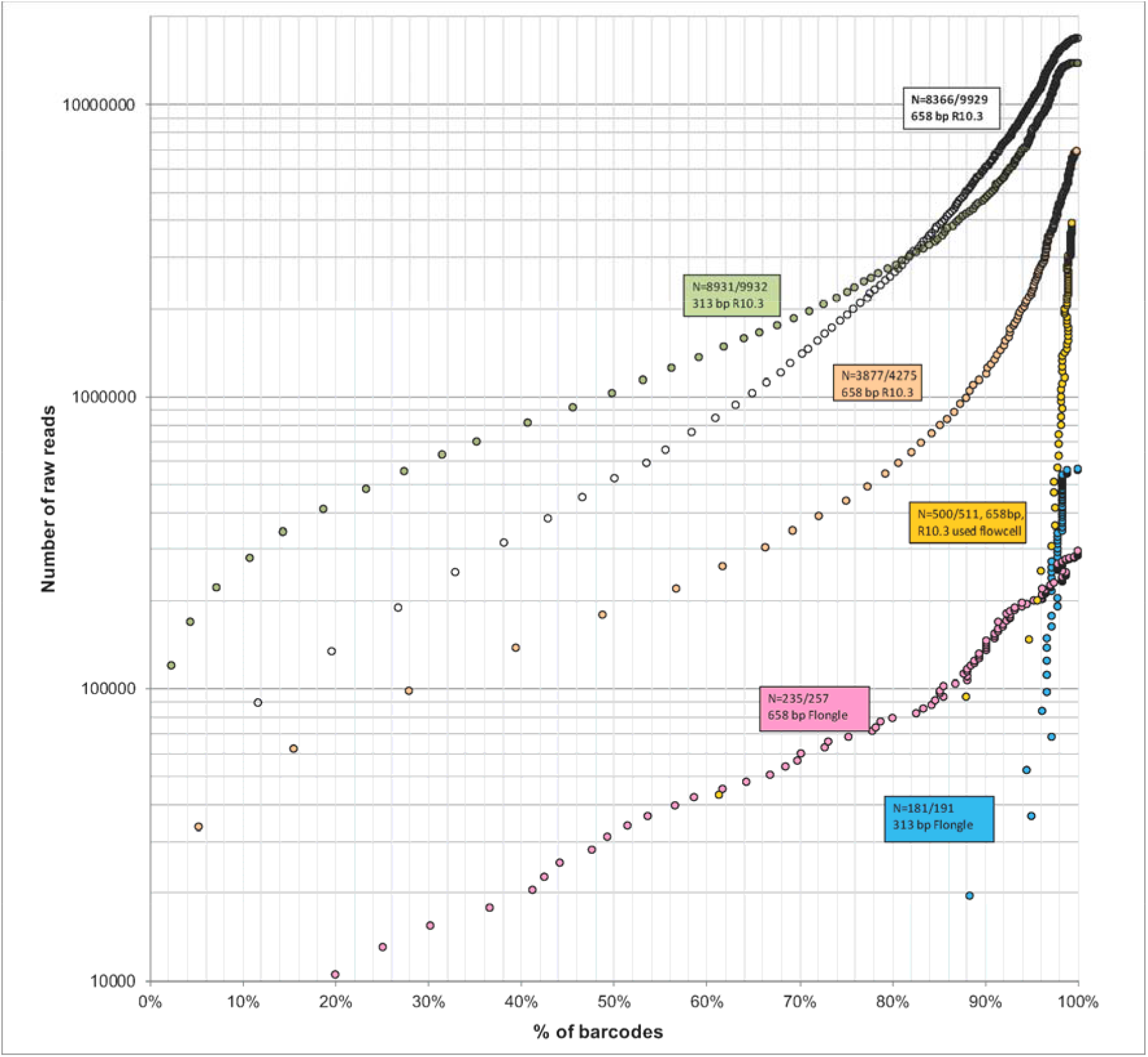
Relationship between barcoding success and number of raw reads for six amplicon pools (191-9932 specimens; barcoding success rates 84-97%). Percentage of barcodes recovered is relative to the final estimate based on all data.

We further compared the barcodes obtained with ONTbarcoder with those obtained via the recent software NGSpeciesID [56]. NGSpeciesID often provides multiple consensus barcodes for the same set of reads for a specimen. We here only compare the consensus barcodes supported by the highest number of reads. When compared to lllumina or Sanger reference barcodes, the barcodes obtained via ONTbarcoder have fewer errors (Table 3: 25-118 erroneous barcodes for NGSpeciesID vs 5-28 erroneous barcodes for ONTbarcoder). In addition, NGSpeciesID also yielded consensus barcodes for negative controls because it performs no quality control based on length, translation or other criteria. Most of the corresponding read sets yielded no barcodes with ONTbarcoder due to rigorous quality checks and low read count filters. For example, NGSpeciesID formed 32 negative consensus barcodes for Afrotropical Phoridae 658 dataset (demultiplexed by minibar) with 30 being represented by <5 sequences.

**Table 3.**
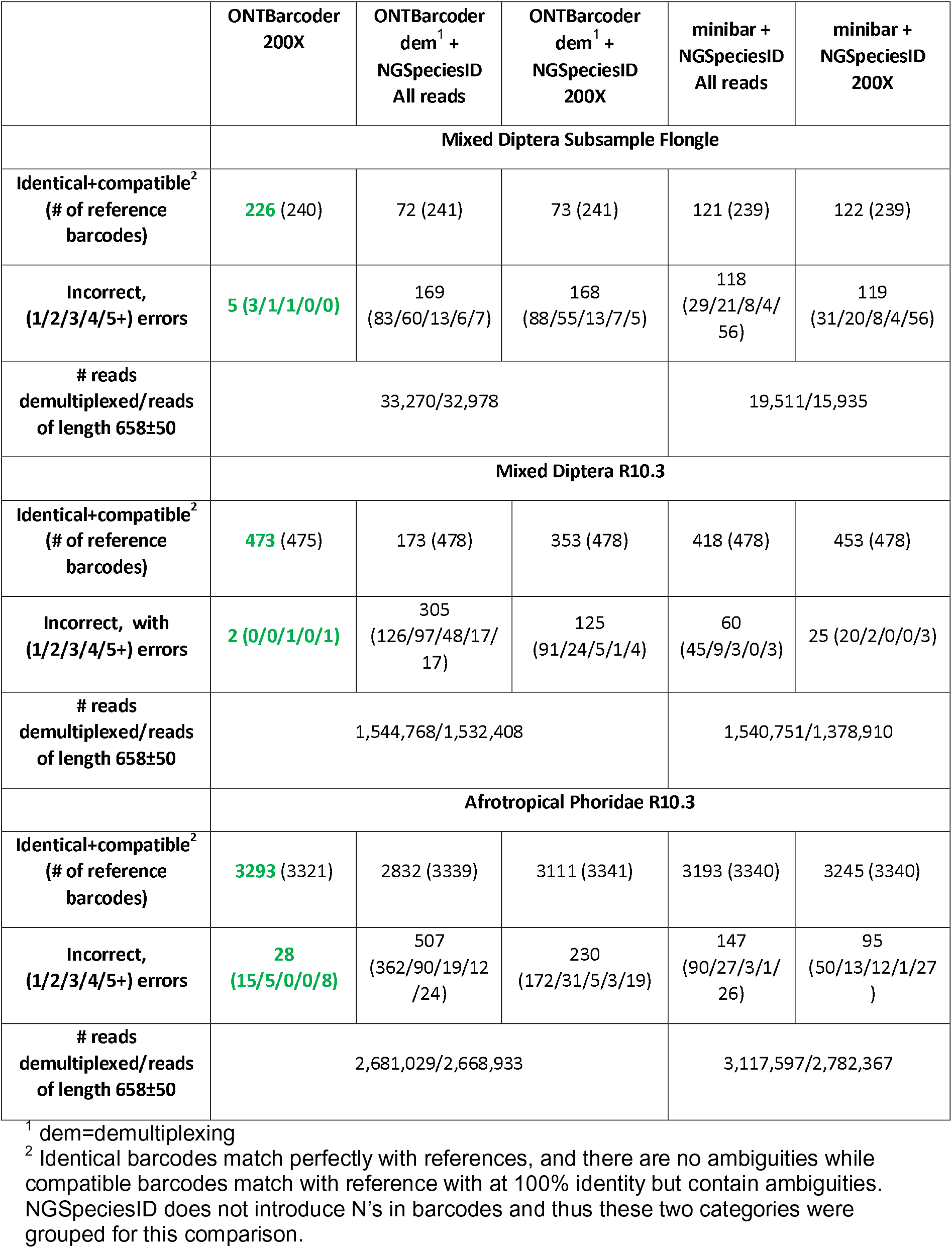
Comparison of barcodes obtained by ONTbarcoder with NGSpeciesID (green=highest number of correct and lowest number of errors for datasets). NGSpeciesID was applied once to the demultiplexed reads obtained with ONTbarcoder and once to those obtained with minibar. Barcode calling was done using all demultiplexed reads as well as 200X subset. ONTbarcoder was run at 200X read coverage (Consensus by Length) and 100X coverage (Consensus by Similarity). The accuracy of MinION barcodes is only compared to reference barcodes obtained with Illumina/Sanger sequencing. Errors are defined as sum of substitution and indel errors.

## Discussion

### Democratization of Barcoding

Biodiversity research needs new scalable techniques for large-scale species discovery and monitoring. This task is particularly urgent and challenging for invertebrates that collectively make up most of the terrestrial animal biomass. We argued earlier that this is likely to be a task that requires the processing of at least 500 million specimens from all over the world with many tropical countries with limited research funding requiring much of the biodiversity discovery work. Pre-sorting these specimens into putative species-level units with DNA sequences is a promising solution as long as obtaining and analyzing the data are sufficiently straightforward and cost-effective. We believe that the techniques described in this manuscript will help with achieving these goals.

We here show that sequencing barcode amplicons with MinION is a particularly attractive option. Firstly, MinION library preparation can be learned within hours and an automated library preparation instrument is available and eventually expected to also generate ligation-based libraries (“VolTRAX”). Secondly, MinION flow cells can accommodate projects of varying scales. Flongle can be used for amplicon pools with a few hundred products, while an R10.3 flow cell can accommodate projects with up to 10,000 specimens. The collection of data on MinION flow cells can be stopped whenever the software controlling the run (“MinKNOW”) indicates that a sufficiently high number of reads have been acquired. Flow cells can then be washed and re-used again although the remaining capacity declines over time so that we have only re-used flow cells up to four times.

Traditionally, the main obstacles to using MinION for barcoding have been poor read quality and high cost. Both issues are fading into the past. The quality of MinION reads has improved to such a degree that the laptop-version of our new software “ONTbarcoder” can generate thousands of very high quality barcodes within hours. There is also no longer a need to “polish” reads or to rely on external data or algorithms. The greater ease with which MinION barcodes can be obtained is due to several factors. Firstly, each flow cell now yields a much larger numbers of reads. Secondly, R10.3 reads have a different error profile which allows for reconstructing higher-quality barcodes. Thirdly, high accuracy basecalling has improved raw read quality and thus demultiplexing rates. Lastly, we can now use parameter settings for MAFFT that are designed for MinION reads. These changes mean that even low-coverage bins can yield very accurate barcodes; i.e., both barcode quality and quantity are greatly improved.

We had previously tested MinION for barcoding in 2018 and 2019 and here re-sequenced some of the same amplicon pools. This allowed for a precise assessment of the improvements. In 2018, sequencing the 511 amplicons of the *Mixed Diptera* sample required one flow cell and we obtained 488 barcodes of which only one lacked ambiguous bases. In 2021, we sequenced on a used flow cell (R10.3) with only ∼500 pores, and we obtained 502 barcodes with >98% (496) being free of ambiguous bases. The results obtained for the 2019 amplicon pools were also better. In 2019, one flow cell (R9.4) allowed us to obtain 3,223 barcodes from a pool of amplicons obtained from 4,275 specimens of *Afrotropical Phoridae*. Resequencing weak amplicons increased the total number of barcodes by approximately 500 to 3,762 [33]. Now, one R10.3 flow cell yielded 3,905 barcodes (+143) for the same amplicon pool, while retaining an accuracy of >99.99% and reducing the ambiguities from 0.45% to 0.01%. If progress continues at this pace, we predict that MinION will be the default barcoding tool for most users. This, too, is because all barcoding steps can now be carried out in one laboratory with a modest set of equipment (see Table 4). With MinION being readily available, there is no longer the need to outsource sequencing and/or to wait until enough barcode amplicons have been prepared for an Illumina or PacBio/Sequel flow cell [57]. This eases biodiversity discovery and allows many biologists, government agencies, students, and citizen scientists from around the globe to get involved in biodiversity discovery.

**Table 4.** Equipment required for MinION barcoding

| Required (<500 specimens) |  |
| --- | --- |
| 1 | MinION sequencer (preferably Mk1C for basecalling) with Flongle adapter |
| 2 | Thermocycler(s) |
| 3 | Gel Electrophoresis setup |
| 4 | Magnetic Separation Rack |
| 5 | Qubit for DNA quantification |
| 6 | Standard equipment: Vortex, Mini-centrifuge, pipettes, freezer, fridge |
| 7 | Standard laptop or PC |
| **Required (>500 specimens)** |  |
| 1 | Multichannel pipette(s) |
| **Optional but highly desirable** |  |
| 1 | Hula Mixer |

This raises the question of how much it costs to sequence a barcode with MinION. There is no straightforward answer because the cost depends on user targets. For example, a user who wants to sequence a pool of 5000 barcodes may want an 80% success rate in order to identify the dominant species in a sample of nuisance midges that occur in mixed-species swarms [58]. Based on Figure 4, only ca. 1.5 million raw MinION reads would be needed. On average, MinION flow cells yield >10 million reads and cost USD 475-900 depending on how many cells are purchased at the same time. Add the library cost of ca. USD 100 and the overall sequencing cost of the project is USD 180-235. This experiment would be expected to yield 4000 barcodes for the 5000 amplicons (4-6 cents/barcode). Given the low cost of 1 million MinION reads ($50-90), we predict that most users will opt for sequencing at a greater depth since this will likely yield several hundred additional barcodes. This will increase the sequencing cost per barcode, because the first 1.5 million reads already recovered barcodes for all strong amplicons. Additional reads will predominantly strengthen read coverage for these amplicons and relatively few reads will be added to the read bins that were too weak to yield barcodes at low coverage; i.e., there are diminishing returns for additional sequencing.

Overall, we thus predict that most users will only multiplex 10,000 amplicons in the same MinION flow cell so that the sequencing cost per specimen would be 0.06-0.10 USD depending on flow cell cost. Large-scale biodiversity projects can reduce the cost further by switching to sequencing with PromethION, a larger ONT sequencing instrument that can accommodate up to 48 flow cells. PromethION flow cells have 6 times the number of pores for twice the cost so that this switch reduces the sequencing cost by 60% (flow cell capacity: 60,000 barcodes). At the other end of the scale are those users who occasionally need a few hundred barcodes. They can use Flongle flow cells, which are comparatively expensive (0.50 USD) because each flow cell costs $70 and requires a library that is prepared with half the normal reagents (ca. $50).

### ONTbarcoder for large-scale species discovery with MinION

We here introduce ONTbarcoder, which runs on a regular laptop using either Windows10, Linux, or Macintosh OS and has a GUI. ONTbarcoder has been extensively tested (>4000 direct comparisons to Sanger and Illumina barcodes) and is designed to yield thousands of barcodes rapidly without impairing accuracy even when encountering low-coverage amplicons. Accuracy is ensured by applying 4 QC criteria related to length and translatability of the barcode. Speed is maintained through the parallelization of most steps on UNIX systems (Mac and Linux; parallelization is restricted to demultiplexing in Windows). ONTbarcoder furthermore allows for updating the parameter file for alignment. This is advisable because MinION continues to evolve quickly. We expect flow cell capacity to increase further and basecalling to improve (see [59]). For example, a new basecaller (“*bonito*”) developed by ONT has shown promise by improving raw read accuracy (https://nanoporetech.com/about-us/news/new-research-algorithms-yield-accuracy-gains-nanopore-sequencing). This basecaller is now also available in MinKNOW and our preliminary tests (Flongle: *Mixed Diptera Subsample, Chironomidae*; R10.3: *Palaearctic Phoridae,* 313 bp; bonito version=0.3.6) confirm that it yields reads of similar quality as HAC (unpublished data). We expect these regular changes to ONT software to further improve the suitability of ONT sequencers for barcoding.

ONTbarcoder evolved from miniBarcoder, which yielded high quality barcodes based on four different tests covering >8000 barcodes [31, 33, 54, 60], but had two drawbacks that have been fixed in ONTbarcoder. Firstly, we dropped the translation-based error correction that tended to increase the number of Ns. This step used to be essential because indel errors were prevalent in consensus barcodes obtained with older flow cell models. Secondly, ONTbarcoder can be installed by unzipping a file and is easy to maintain on different operating systems. Until now, external dependencies meant that several software packages had to be installed and that only some operating systems were compatible. This has been a major drawback of all MinION bioinformatics pipelines and led Watsa et al. [35] to recommend that bioinformatics training is needed before MinION barcoding could be used in schools (e.g., training in UNIX command-line). With ONTbarcoder, such training will no longer be needed.

MinION has been used for barcoding fungi, animals, and plants and alternative pipelines have been developed [36, 54, 56, 60–66], but there is one fundamental difference between these studies/pipelines and the vision presented here. These studies tended to show that MinION sequencing can be done in the field. Thus only a very small number of specimens were analysed (<150 with the exception of >500 in Chang, Ip et al [60]). The potential use in the field is an attractive feature for time-sensitive samples that could degrade before reaching a lab. However, it is unlikely to help substantially with tackling large-scale biodiversity discovery and monitoring because obtaining few MinION barcodes per flow cell is too expensive. Additionally, the bioinformatic pipelines that were developed for these small-scale projects were not suitable for large-scale, decentralized barcoding. For example, some used ONT’s commercial barcoding kit that only allows for multiplexing up to 96 samples in one flow cell [64, 66]; i.e., each amplicon has very high read coverage which influenced the design of the analysis pipelines (e.g. coverage recommendations for ONTrack is 1000x: Maestri, Cosentino et al.[64]). The high coverage requirements also meant that the pipelines were only tested for small numbers of samples (<60: Menegon, Cantaloni et al. 2017, Maestri, Cosetino et al. 2019, Seah, Lim et al. 2020, Sahlin, Lim et al. 2021 [56, 61, 64, 66]) which are unlikely to represent the complexities of large, multiplexed amplicon pools (e.g., nucleotide diversity, uneven coverage).

This concern is confirmed by our test of the most recently introduced bioinformatics pipeline (NGSpeciesID [56]). It requires minibar/qcat and nanofilt, isONclust SPOA, Parasail, and optionally, Medaka [63, 67, 68] and often yields multiple consensus barcodes for the same set of reads because it relies on an intermediate clustering step. To assess the performance of NGSpeciesID, we used the consensus barcode with the highest coverage and only compared the results for those barcodes for which we had reference barcodes obtained with Sanger and Illumina (Table 3). We find that under optimal settings, NGSpeciesID yields 3-23 times the number of erroneous barcodes than ONTbarcoder (Table 3). One reason is that NGSpeciesID does not use ambiguity codes in consensus sequences; i.e., ONTbarcoder will place an “N” when the evidence is ambiguous while NGSpeciesID will opt for one of the nucleotides. In addition, NGSpeciesID does not use barcode length or translatability as QC criteria. This has the downside that the software is more likely to yield erroneous barcodes. For example, NGSpeciesID proposes consensus barcodes for very small read sets for which ONTbarcoder proposes no barcodes because they failed the QC or did not meet the minimum read threshold. An upside of not using barcode length and translatability as QC is that NGSpeciesID can propose consensus barcodes for genes that are non-coding (e.g., ITS) or have high length variability (e.g., ribosomal genes). However, the user should be aware that 3-5% of these barcodes will include errors if the results for *COI* also hold for other genes (see Table 3).

### Biodiversity monitoring with MinION barcodes

Despite the widespread use of metabarcoding for analyzing samples consisting of hundreds or thousands of specimens [69, 70], large-scale barcoding of individual specimens remains essential for discovering and describing species. It associate barcodes with individual voucher specimens, which can be used for further research. This is essential for taxonomic research, which is the only way to fix systematic errors caused by *COI* which lumps recently diverged species and splits species with deep allopatric splits [71]. High quality barcode databases are also important for the analysis of metabarcoding data because they facilitate the identification of numts, heteroplasmy, contaminants and errors. Furthermore, large-scale barcoding will also be critical for developing AI-assisted biodiversity monitoring of invertebrates using images [72]. Such monitoring requires neural networks that are trained with large numbers of images. These images are best obtained from specimens that were identified/grouped into species based on barcodes barcodes. It appears likely that AI-assisted biodiversity monitoring will be the method of choice in the future because it can has the potential to quickly identify and count common species and highlight which specimens may belong to new/rare species [73].

## Conclusions

Many biologists would like to have ready access to barcodes without having to send specimens halfway around the world or run large, complex, and expensive molecular laboratories. Many have been impressed by MinION’s low cost, portability, and its ability to deliver real-time sequencing, but they were worried about high cost and complicated bioinformatics pipelines. We here demonstrate that these concerns are no longer justified. MinION barcodes obtained by R10.3 flow cells are virtually identical to barcodes obtained with Sanger and Illumina sequencing. Barcoding with MinION is now also cost-effective and the new “ONTbarcoder” software makes it straightforward to analyze the data on a standard laptop. Add the simplified and cheaper methods for obtaining amplicons and biodiversity discovery will become more scalable and accessible to all.

## Methods

MinION and Flongle sequencing were here tested for six amplicon pools (Table 5). For two of the pools, *Mixed Diptera* (N=511) and *Afrotropical Phoridae* (N=4,275), we already had amplicons and comparison barcodes that were obtained with Sanger, Illumina, and older versions of MinION flow cells that used a different chemistry (see below). These two pools were here used to assess the accuracy of barcodes generated using the new MinION flow cell using the R10.3 chemistry. Two additional datasets were used to test the capacity of R10.3 flowcells for mini- and full length barcodes by sequencing barcodes of different lengths for the same specimens and obtained with the same DNA template (*Palaearctic Phoridae,* 658 and 313 bp for ca. 9,930 specimens). Lastly, we tested the performance of Flongle flow cells using a *Chironomidae* dataset (313 bp mini-barcodes for 191 specimens) and a *Mixed Diptera Subsample* (full-length barcodes for 257 specimens) of the aforementioned *Mixed Diptera* amplicon pool for which we had Sanger barcodes for comparison.

**Table 5.** Datasets used in the study and the corresponding experimental details.

| <u>Dataset Name</u> | <u>Number of specimens</u> | <u>Fragment size, primer information</u> | <u>Extraction/PCR setup</u> | <u>PCR cleanup</u> | <u>ONT Library Preparation kit/Flow cell used</u> |
| --- | --- | --- | --- | --- | --- |
| **R10.3 Datasets** |  |  |  |  |  |
| <u>Mixed Diptera</u> | 511 (257 mixed | 658 bp | Extraction Method: | Ampure | SQK- |
| (see Srivathsan et al. 2018)<br>- Sanger barcodes available | Diptera, 254 Dolichopodidae)<br>17 negatives | HCO2198, LCO1490 [75] | QuickExtract PCR Mix:<br>Total volume: 20 µl<br>10x buffer: 2 µl<br>dNTPs (2.5 mM): 1.5 µl<br>Taq polymerase: 0.2 µl<br>BSA (1 mg/ml): 2 µl<br>Primer (5 µM): 2 µl each<br>DNA: 2 µl | beads (Beckman Coulter) | LSK110/FLO-MIN111 |
| <u>Afrotropical Phoridae</u> (see Srivathsan et al. 2019)<br>- Illumina mini-barcodes available | 4275 (Phoridae)<br>45 negatives | 658 bp HCO2198, LCO1490 [75] | Extraction Method: QuickExtract PCR Mix:<br>Total volume: 15.16 µl<br>Mastermix (CWBio): 10 µl<br>25mM MgCl<sub>2</sub>: 0.16 µl<br>BSA (1mg/ml): 2 µl<br>Primer (10µM): 1 µl each<br>DNA: 1 µl | Sera-Mag beads (GE Healthcare Life Sciences) in PEG | SQK-LSK109/FLO-MIN111 |
| <u>Palaeartic Phoridae (658)</u> | 9,929 (Phoridae)<br>105 negatives | 658 bp jgHCO2198, LCO1490 [75, 76] | Extraction Method: HotSHOT PCR Mix:<br>Total volume: 16 µl<br>Mastermix (CWBio): 7 µl<br>BSA (1mg/ml): 1 µl<br>Primer (10µM): 1 µl each<br>DNA: 6 µl | Ampure beads (Beckman Coulter) | SQK-LSK110/FLO-MIN111 |
| <u>Palaeartic Phoridae (313)</u> | 9,932 (Phoridae)<br>106 negatives | 313 bp m1COLintF, jgHCO2198 [74, 76] | Extraction Method: HotSHOT PCR Mix:<br>Total volume: 14 µl<br>Mastermix (CWBio): 7 µl<br>BSA (1mg/ml): 1 µl<br>Primer (10µM): 1 µl each<br>DNA: 4 µl | Ampure beads (Beckman Coulter) | SQK-LSK110/FLO-MIN111 |
| **Flongle Datasets** |  |  |  |  |  |
| <u>Mixed Diptera subsample</u> (see Srivathsan et al. 2018)<br>- Sanger barcodes available | 257<br>7 negatives | See "Mixed Diptera" entry for R10.3 | See "Mixed Diptera" entry for R10.3 | Ampure beads (Beckman Coulter) | SQK-LSK109/Flongle |
| <u>Chironomidae</u> | 191 (Chironomidae)<br>1 negative | 313 bp m1COLintF, jgHCO2198 [74,76] | Extraction Method: HotSHOT PCR Mix:<br>Total volume: 14 µl<br>Mastermix (CWBio): 7 µl<br>BSA (1mg/ml): 1 µl<br>Primer (10µM): 1 µl each<br>DNA: 4 µl | Ampure beads (Beckman Coulter) | SQK-LSK109/Flongle |

The methods for obtaining the reference barcodes with Sanger and Illumina are described in Srivathsan et al. [31, 33]. Briefly, for Sanger sequencing (Mixed Diptera sample set), the same PCR products sequenced with MinION were individually cleaned using SureClean Plus (Bioline, London) and subjected to cycle sequencing using BigDye^TM^. The products were precipitated using PureSeq (Aline BioSciences, Woburn) and analysed in ABI 3730xl 96 capillary sequences. The resulting chromatograms were edited using Sequencher v4 (GeneCodes, Ann Arbor). For Illumina sequencing of products of the Afrotropical Phoridae dataset, an independent PCR was conducted using the DNA extract for the same specimens to amplify a short 313-bp fragment of COI using 9-bp tagged versions of the primers described by Leray et al. [74]. The products were pooled and sequenced using HiSeq2500 (250 bp PE sequencing). The data processing followed Wang et al.’s protocol. [23] for obtaining a set of consensus barcodes.

## 1: DNA extraction

For all newly barcoded specimens, we used DNA template obtained with 10-15 μL HotSHOT per specimen [77], but other buffers like PBS could have also been used (see [78]). Small specimens were submerged within the well of a microplate while larger specimens were placed head-first into the well. Note that the specimen need not be entirely submerged in HotSHOT. The placement of 95 specimens takes approximately 17 minutes and the DNA is obtained within 20 minutes in a thermocycler via two heating steps [77] (https://www.youtube.com/watch?v=y1qGzL5PraQ&;t=3s). After neutralization, >20 l of μl of template is available for amplifying *COI* and the voucher can be recovered. The advantages of HotSHOT is low cost and speed. The disadvantages are fast degeneration of the leftover template within days. Some alternatives to DNA extraction with HotSHOT are described in Table 6. We consider them too costly or time-consuming given that *COI* is a mitochondrial gene and thus naturally enriched. Indeed, the small mitochondrial genome (16 kbp) usually contributes 0.5-5% of the DNA in a genomic extraction [79, 80]. Therefore obtaining sufficient template for DNA barcoding should take <20 minutes, does not require DNA purification, and should cost essentially nothing as long as the specimens contain DNA of reasonable quality (e.g., <20 year old Malaise trap samples).

**Table 6.**
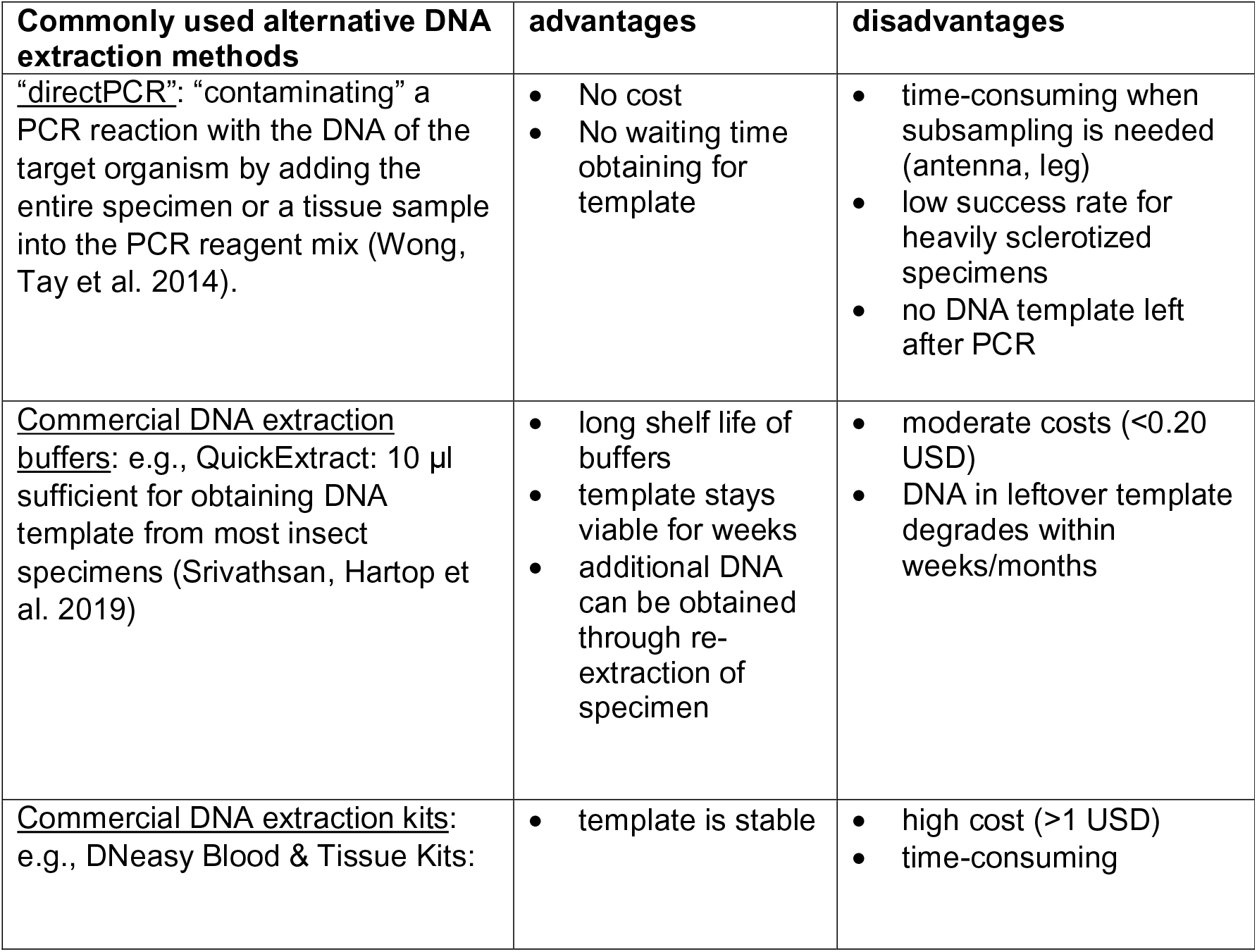
Alternative DNA extraction methods

| Commonly used alternative DNA extraction methods | advantages | disadvantages |
| --- | --- | --- |
| <u>“directPCR”</u>: “contaminating” a PCR reaction with the DNA of the target organism by adding the entire specimen or a tissue sample into the PCR reagent mix (Wong, Tay et al. 2014). | No cost<br>No waiting time obtaining for template | time-consuming when subsampling is needed (antenna, leg)<br>low success rate for heavily sclerotized specimens<br>no DNA template left after PCR |
| <u>Commercial DNA extraction buffers</u>: e.g., QuickExtract: 10 µl sufficient for obtaining DNA template from most insect specimens (Srivathsan, Hartop et al. 2019) | long shelf life of buffers<br>template stays viable for weeks<br>additional DNA can be obtained through re-extraction of specimen | moderate costs (<0.20 USD)<br>DNA in leftover template degrades within weeks/months |
| <u>Commercial DNA extraction kits</u>: e.g., DNeasy Blood & Tissue Kits: | template is stable | high cost (>1 USD)<br>time-consuming |

## 2: Amplification of COI via PCR

Obtaining amplicons for DNA barcodes does not require high-fidelity polymerases, which are mostly needed for amplifying low copy-number nuclear genes based on low-concentration template. Standard polymerases are sufficient. We exclusively used CWBio 2x master mix in 14-16 ul PCR reactions (see Table 5 for details). The set-up of one plate requires <15 minutes (https://www.youtube.com/watch?v=NxYOvZGhD0E&t=5s). In order to save time, only a small number of reactions per microplate need to be checked via gel electrophoresis (N=8-12, including the negative control). Running out additional amplicons is not necessary because failed amplicons do not add to the MinION sequencing cost and can be recovered via re-sequencing [33] or re-amplification.

Amplicon sequencing with all second and third generation sequencing technologies (including MinION) involves amplicon pools. This means that the amplicon for each specimen has to be tagged/indexed/barcoded with short DNA sequences at the 5’ ends of the amplicons. This allows for the assignment of each read obtained during sequencing to a specific specimen during the “demultiplexing” bioinformatics step. We use 13 bp tags that are distinct (>4bp from each other including insertions/deletions) and lack homopolymers (see supplementary materials). This tag length is a compromise given that longer tags have the disadvantage of reducing PCR success rates [33] while having the advantage of increasing the proportion of reads that can be demultiplexed with confidence.

Numerous dual-PCR tagging techniques for amplicons have been described in the literature [29, 64, 81, 82], but we only use single-PCR tagging [28]. It is here described for a microplate with 96 templates, but the same principle can be applied to strip tubes or partial microplates. What is needed is a 96-well primer plate where each well contains a differently tagged reverse primer. This “primer plate” can yield 96 unique combinations of primers once the 96 reverse primers are combined with one forward primer (f-primer x 96 differently tagged r-primers = 96 unique combinations). This also means that if one purchases 105 differently tagged forward primers, one can individually tag 10,800 specimens (105 x 96= 10,800 amplicons). This is the number of amplicons that we consider appropriate for a MinION flow cell (R10.3; see Results). In the laboratory we assign the tag combinations as follows. For each plate with 96 PCR reactions, we add the same f-primer to a tube with the PCR master mix to be used for the entire plate. We then dispense the “f-primed” master mix into the 96-wells. Afterwards, we use a multichannel pipette to add the DNA template and the tagged r-primers from the r-primer plate into the PCR plate. All 96 samples in the plate now have a unique combination of tagged primers because they only share the same tagged forward primer. This makes tracking of tag combinations simple because each PCR plate has its own tagged f-primer, while the r-primer is consistently tied to well position. Each plate has a negative control that is used to detect contamination. The tagging information for each plate is recorded in the demultiplexing file that is later used to demultiplex the reads obtained during sequencing (see supplement for an example).

We prefer single-PCR tagging over two-PCR tagging [29, 64, 81, 82] because it is cheaper (requires half the amount of primer), less error-prone (fewer PCR reactions and cycles), and saves time (no need to clean-up the first-round amplicons). The only downsides of single-PCR tagging are an initially higher investment in primers and the need to manage the primer stock more carefully because it is used for a longer time. Long-term storage should thus be at -80°C and the number of freeze-thaw cycles should be kept low (<10).

## 3. Amplicon sequencing

PCR is followed by pooling, purification of the amplicons via the removal of unused PCR reagents, the adjustment of DNA concentration, and sequencing. We only pool 1 μl of each PCR product. The pools were cleaned using SPRI bead-based clean-up with Ampure (Beckman Coulter) beads, but Kapa beads (Roche) or the more cost-effective Sera-Mag beads (GE Healthcare Life Sciences) in PEG [83] are also viable options [33]. For barcodes longer than 300 bp, we recommend the use of a 0.5X ratio for Ampure beads since it removes a larger proportion of primers and primer dimers. However, this ratio is only suitable if yield is not a concern (e.g., pools consisting of many and/or high concentration amplicons). Increasing the ratio to 0.7-1X will improve yield but render the clean-up less effective. The pooling and clean-up of three 96-well plates takes about 40 minutes (https://www.youtube.com/watch?v=YKVWEvcSw6A), but the time per plate is lower when large numbers of amplicons are pooled. Amplicon pools containing large numbers of amplicons may require multiple rounds of clean-up, but only a small subset of the initial pool has to be purified because most library preparation kits require only small amounts of DNA. We confirm the success of the clean-up procedures via gel electrophoresis, which should show only one strong band of expected length. After the clean-up, the pooled DNA concentration is measured in order to use an appropriate amount of DNA for library preparation. We use Qubit, but less precise techniques are probably also suitable.

Obtaining a cleaned amplicon pool according to the outlined protocol is not time consuming, but many studies retain “old Sanger sequencing habits”. For example, they use gel electrophoresis for each PCR reaction to test whether an amplicon has been obtained. Afterwards, they clean and measure all amplicons – one at a time – for normalization (often with very expensive techniques: Ampure beads: [64]; TapeStation, BioAnalyzer, Qubit:[66]). This is presumably done to obtain a pool of amplicons where each has equal representation. However, reads are cheap while the clean-up and measurement of many amplicons is expensive and unnecessary because weak products that failed to yield a barcode in the first sequencing run can be re-sequenced [33].

## 4. Library preparation and sequencing

We prepare our MinION libraries using ligation-based kits and 200 ng of DNA for full flow cells and 100 ng for Flongle (see Table 6 for details). We generally follow kit instructions, but exclude the FFPE DNA repair mix in the end-repair reaction, as this is mostly needed for formalin-fixed, paraffin-embedded samples. The reaction volumes for the R10.3 flow cell libraries consist of 45 μl of DNA, 7 μl of Ultra II End-prep reaction buffer (New England Biolabs), 3 μl of Ultra II End Prep enzyme mix (New England Biolabs) and 5 μl of molecular grade water. For the Flongle, only half of the reagents are used to obtain a total volume of 30 μl. We further modify the Ampure ratio to 1x for all steps as DNA barcodes are short whereas the recommended ratio in the manual is for longer DNA fragments. The libraries for the experiments in this study were loaded and sequenced on a MinION Mk 1B. Data capture involved a MinIT or a Macintosh computer that meets the IT specifications recommended by ONT. The bases were called using Guppy (versions provided in Table 1), under the high616 accuracy model in MinIT taking advantage of its GPU.

We here only used MinION and Flongle flow cells for barcoding. This preference was based on four considerations: (1) Scaling; i.e., ability to accommodate projects of different sizes, (2) turnaround times, (3) cost, and (4) amplicon length. Flongle can be used for small amplicon pools (<300 products) because it has low fixed costs per experiment (library and flow cell: ca. $120 USD) and the turnaround time is fast, so the MinION Flongle is arguably the best sequencing option for small barcoding projects with > 50 barcodes. Full MinION flow cells also have fast turnaround times, but the minimum run cost is closer to 1000 USD for most users (flow cell cost drops with bulk purchase), so this option only becomes more cost-effective than Flongle when >1800 amplicons are sequenced. As illustrated in the Results section of the manuscript, one MinION flow cell can comfortably sequence 10,000 amplicons.

Alternatives to MinION/Flongle are sequencing barcode amplicons with Sanger, Illumina (Wang, Srivathsan et al. 2018), or PacBio (e.g., Sequel: Hebert, Braukmann et al. [29]). Sanger sequencing has fast turnaround times but high sequencing cost per amplicon ($3-4 USD). PacBio’s Sequel flow cells have a similar capacities as full MinION flow cells [29] and the consumable costs are also similar for most users who have to outsource Sequel sequencing due to the high instrument cost for PacBio. However, Sequel does not allow for flexible scaling like Flongle/MinION and most users will have to wait several weeks until the data are returned from the service provider. By far the most cost-effective sequencing method for barcodes is Illumina’s NovaSeq sequencing. However, the fixed costs for library and lanes are high (3000-4000 USD), but each flow cell yields 800 million reads which can comfortably sequence 800,000 barcodes at a cost of < $0.01 USD per barcode. Note that Illumina reads are only suitable for generating mini-barcodes of up to 420 bp length (using 250bp PE sequencing using SP flow cell). “Full-length” *COI* barcode (658 bp) can only be obtained by sequencing two amplicons per specimen.

## 5. Bioinformatics: Development and application of ONTbarcoder

One of the most significant barriers to widespread barcoding with MinION has been complex bioinformatics pipelines that were needed for fixing the high error rates of ONT reads. However, after obtaining data from a new R10.3 flow cell, we noticed major improvements in read quality, the total number of raw reads, and the number of demultiplexed reads. This led to the development of “ONTbarcoder”, which has a graphical user interface (GUI) and is suitable for all major operating systems (Linux, Mac OS, Windows10). The use of the software is illustrated in a video tutorial: https://www.youtube.com/channel/UC1WowokomhQJRc71FmsUAcg.

### ONTbarcoder

ONTbarcoder (available at: https://github.com/asrivathsan/ONTbarcoder) has three modules. (a) The first is a demultiplexing module which assigns reads to specimen-specific bins. (b) The second is a barcode calling module which reconstructs the barcodes based on the reads in each specimen bin. (c) The third is a barcode comparison module that allows for comparing barcodes obtained via different software and software settings.

#### a. Demultiplexing

In order to obtain barcodes three pieces of information and two files have to be provided to ONTbarcoder via the GUI: (1) primer sequence, (2) expected fragment length, and (3) demultiplexing information (=tag combination for each specimen). The latter is summarized in a demultiplexing file (see supplementary information for format). The only other required file is the FASTQ file obtained from MinKNOW/Guppy after basecalling. Demultiplexing by ONTbarcoder starts by analyzing the read length distribution in the FASTQ file. Only those reads that meet the read length threshold are demultiplexed (default= 658 bp corresponding to metazoan COI barcode). Technically, the threshold should be the amplicon length plus the length of both tagged primers, but ONT reads have indel errors such that they are occasionally too short or too long. We therefore advise to specify the amplicon length without primer&tag as threshold. ONTbarcoder will split reads that are twice the expected fragment length into two parts whose lengths are determined based fragment size, primer and tag lengths, and a window to account for indel errors (default=100 bp).

Once all reads with a suitable length for demultiplexing have been identified, ONTbarcoder finds the primers via sequence alignment of the primer sequence to the reads (using python library *edlib*). Up to 10 deviations from the primer sequence are allowed because this step is only needed for determining the primer location and orientation within the read. For demultiplexing based on the tags, the flanking region of the primer sequence is retrieved whereby the number of retrieved bases is equal to the user-specified tag length. The flanking sequences are then matched against the tags from the user-provided tag combinations (demultiplexing file). In order to account for sequencing errors, not only exact matches are accepted, but also matches that differ by up to 2 bps from the tag sequence (substitutions/insertions/deletions). Note that accepting tag variants does not lead to demultiplexing error because all tags differ by >4 bp. All reads with the same tag combination thus identified belong to the same specimen and are pooled into the same bin. To increase efficiency, demultiplexing is parallelized and the search space for primers and tags are restricted to the beginning and end of the reads (window is user-specified).

#### b. Barcode calling

Barcode calling uses the reads within each specimen-specific bin to reconstruct the barcode sequence. The reads are aligned to each other and a consensus sequence is called. Barcode calling is done in three phases: “Consensus by Length”, “Consensus by Similarity” and “Consensus by barcode comparison”. The user can opt to only use some of these methods.

“Consensus by Length” is the main barcode calling mode and relies on efficient alignment in order to provide reasonable speed for thousands bins each containing many reads. ONTbarcoder delivers speed by using an iterative approach that gradually increases the number of reads (“coverage”) per bin that is used during alignment. However, reconstructing barcodes based on few reads could lead to errors which are weeded out by ONTbarcoder by applying four Quality Control (QC) criteria. The first three QC criteria are applied immediately after the consensus sequence has been called: (1) the barcode must be translatable, (2) it has to match the user-specified barcode length, and (3) the barcode has to be free of ambiguous bases (“N”). To increase the chance of finding a barcode that meets all three criteria, we subsample the reads in each bin by read length (thus the name “Consensus by Length”); i.e., initially only those reads closest to the expected length of the barcode are used. For example, if the user specified coverage=25x for a 658 bp barcode, ONTbarcoder would only use the 25 reads that have the closest match to 658 bp. The fourth QC measure is only applied to barcodes that have already met the first three QC criteria. A multiple sequence alignment (MSA) is built for the barcodes obtained from the amplicon pool, and any barcode that causes the insertion of gaps in the MSA is rejected. Note that if the user suspects that barcodes of different length are in the amplicon pool, the initial analysis should use the dominant barcode length. The remaining barcodes can then be recovered by re-analyzing all data or only the failed read bins (“remaining”, see below) and bins that yielded barcodes that had to be “fixed”. These bins can be reanalyzed using a different pre-set barcode length.

“Consensus by Similarity”. Barcodes that failed the QC during the “Consensus by Length” stage are often very close to the expected length and have few ambiguous bases, and/or cause few gaps in the MSA. These “preliminary barcodes” can be improved through “Consensus by Similarity”. This method eliminates outlier reads from the read pool in the bins. Such reads can differ considerably (see below) from the signal of the consensus barcode and ONTbarcoder identifies them by sorting all reads by similarity to the preliminary barcode. Only the top 100 reads (this default can be changed) that differ by <10% from the preliminary barcode are retained and used for calling the barcodes again using the same techniques described “Consensus by Length” (including the same QC criteria). This improvement step converts many preliminary barcodes found during “Consensus by Length” into barcodes that pass all four QC criteria by filling/removing indels or resolving an ambiguous base.

“Consensus by barcode comparison”. The remaining preliminary barcodes that still failed to convert into QC-compliant barcodes tend to be based on read bins with low coverage, but some can yield good barcodes after subjecting them to a further improvement step that fixes the remaining errors. ONTbarcoder identifies these errors by finding the 20 most similar QC-compliant barcodes that have already been reconstructed for the other amplicons. The 21 sequences are aligned and ONTbarcoder finds the errors because they cause insertions and deletions in the MSA. Insertions are deleted, gaps are filled with ambiguous bases (“N”), but mismatches are retained. The number and kinds of “fixes” are recorded and added to the FASTA header of the barcode. Large number of fixes imply that the barcode should not be used (see below). “Rare taxa are disadvantaged by this method, but the barcode for very few if any will ever reach the “Consensus by barcode comparison” stage because most/all will be resolved earlier. We added “consensus by barcode comparison” because it helps with resolving weak barcodes for specimens that represent abundant species.

Output. ONTbarcoder produces a summary table (Outputtable.csv) and FASTA files that contain the different classes of barcodes. Each barcode header contains information on coverage used for barcode calling, coverage of the specimen bin, length of the barcode, number of ambiguities and number of indels fixed. Five sets of barcodes are provided, here discussed in the order of barcode quality: (1) “QC_compliant”: The barcodes in this set satisfy all four QC criteria without correction and are the highest quality barcodes. (2) “Filtered_barcodes”: this file contains the barcodes that are translatable, have <1% ambiguities and have up to 5 indels fixed during the last step of the bioinformatics pipeline. These filtering thresholds were calibrated based on two datasets for which we have Sanger/Illumina barcodes and the resulting barcodes are found to be highly accurate. Note that the file with filtered barcodes also includes the QC_compliant barcodes and that all results discussed in this manuscript are based on filtered barcodes given that they are of much higher quality than the average barcode in BOLDSystems (assessment in Srivathsan, Balğlu et al [31]).

The remaining files include barcodes of lesser and/or suspect quality. (3) “Fixed_barcodes_XtoY”: these files contain barcodes that had indel errors fixed and are grouped by the number of errors fixed. Only the barcodes with 1-5 errors overlap with Filtered barcodes file, if they have <1% ambiguities. (4) “Allbarcodes”: this file contains all barcodes in sets (1)-(3). (5) “Remaining”: these are barcodes that fail to either translate or are not of predicted length. Note that all barcodes should be checked via BLAST against comprehensive databases in order to detect lab contamination. There are several online tools available for this and we recommend the use of GBIF sequence ID tool (https://www.gbif.org/tools/sequence-id) which gives straightforward output including a taxonomic summary.

The output folder also includes the FASTA files that were used for alignment and barcode calling. The raw read bins are in the “demultiplexed” folder, while the resampled bins (by length, coverage, and similarity) are in their respective subfolders named after the search step. Note that the raw reads are encoded to contain information on the orientation of the sequence and thus cannot be directly used in other software without modifications (see ONTbarcoder manual on Github). Lastly, for each barcode FASTA file (1-5), there are folders with the files that were used to call the barcodes. This means that the user can, for example, reanalyze those bins that yielded barcodes with high numbers of ambiguous bases. Lastly a “runsummary.xlsx” document allows the user to explore the details of the barcodes obtained at every step of the pipeline.

### Algorithms

ONTbarcoder uses the following published algorithms. All alignments utilize MAFFTv7 (Katoh and Standley 2013). The MinION reads are aligned using an approach similar to lamassemble [84] with parameters optimized for nanopore data by “last-train” [85] which accounts for strand specific error biases. The MAFFT parameters can be modified in the “parfile” supplied with the software which helps with adjusting the values given the rapidly changing nanopore technology. All remaining MSAs in the pipeline (e.g., of preliminary barcodes) use MAFFT’s default settings. All read and sequence similarities are determined with the *edlib* python library under the Needle-Wunsch (“NW”) setting, while primer search is using the infix options (“HW”). All consensus sequences are called from within the software. This is initially done based on a minimum frequency of 0.3 for each position. This threshold was empirically determined based on datasets where MinION barcodes can be compared to Sanger/Illumina barcodes. The threshold is applied as follows. All sites where >70% of the reads have a gap are deleted. For the remaining sites, ONTbarcoder accepts those consensus bases that are found in at least >30% of the reads. If no base/multiple bases reach this threshold, an “N” is inserted. To avoid reliance on a single threshold, ONTbarcoder allows the user to change the consensus calling threshold from 0.2 to 0.5 for all barcodes that fail the QC criteria at 0.3 frequency. However, barcodes called at different frequencies are only accepted if they pass the first three QC criteria and are identical. If no such barcode is found, the 0.3 frequency consensus barcode is used for further processing.

#### c. Barcode comparison

Many users may want to call their barcodes under different settings and then compare barcode sets. The ONTbarcoder GUI simplifies such comparisons. A set of barcodes can be dragged into the window and the user can select a barcode set as the reference. The barcode comparisons are conducted using *edlib* library. The barcodes in the sets are compared and classified into three categories: “identical” where sequences are a perfect match and lack ambiguities, “compatible” where the sequences only differ by ambiguities, and “incorrect” where the sequences differ by at least one base pair. Several output files are provided. A summary sheet, a FASTA file each for “identical”, “compatible”, and the sequences only found in one dataset. Lastly, there is a folder with FASTA files containing the different barcodes for each incompatible set of sequences. This module can be used for either comparing set(s) of barcodes to reference sequences, or for comparing barcode sets against each other. It furthermore allows for pairwise comparisons and comparisons of multiple sets in an all-vs-all manner. This module was here used to get the final accuracy values presented in Table 2.

### Quality of Flongle and MinION barcodes

We first used ONTbarcoder to analyze the data for all six datasets by analyzing all specimen-specific read bins at different coverages (5-200x in steps of 5x). This means that the barcodes for a bin with 27 reads were called five times at 5x, 10x, 15x, 20x, and 25x coverages while bins with >200x were analyzed 40 times at 5x increments. Instead of using conventional rarefaction via random subsampling reads, we used the first reads provided by the flow cell because this accurately reflects how the data accumulated during the sequencing run and how many barcodes would have been obtained if the run had been stopped early. This rarefaction approach also allowed for mapping the barcode success rates against either coverage or time.

In order to obtain a “best” estimate for how many barcodes can be obtained, we also carried out one analysis at 200x coverage with the maximum number of “Comparison by Similarity” reads set to 100. All analyses produced a “filtered” set of barcodes (barcodes with <1% Ns and up to 5 fixes) that were used for assessing the accuracy and quality via comparison with Sanger and Illumina barcodes for *Mixed Diptera (MinION R10.3)*, *Afrotropical Phoridae (MinION R10.3)*, and *Mixed Diptera Subsample (Flongle R9.4).* For the comparisons of the barcode sets obtained at the various coverages, we used MAFFT and the assess_corrected_barcode.py script in miniBarcoder (Srivathsan et al. 2019).

## 6. Bioinformatics: Application of minibar and NGSpeciesID

We compared the barcodes obtained with ONTbarcoder with those obtained with the recently published NGSpeciesID [56]. This comparison was carried out for the datasets that have the reference barcodes obtained with Sanger and Illumina (*Mixed Diptera (MinION R10.3)*, *Afrotropical Phoridae (MinION R10.3)*, and *Mixed Diptera Subsample (Flongle R9.4)*). Two comparisons were made. Firstly, we used the same demultiplexed reads that were used for calling the barcodes using ONTbarcoder. Secondly, we demultiplexed the data using *minibar* (git commit: 938ae51) and applied NGSpeciesID (Git commit: 24afc6c) for consensus barcode calling. This approach allowed for software comparisons for consensus calling and demultiplexing. *minibar* was run using the parameters (-e 2 -E 10 -T), i.e. maximum number of errors in tag region was set to 2, in primer alignment was set to 10 and primer and tags were trimmed from the sequences. NGSpeciesID was run with both full datasets as well as by subsampling the data to 200 reads (--sample_size 200) to keep it comparable with our analysis using ONTbarcoder. Other parameters settings were: intended target length = 658 (--m) and maximum deviation from target length = 50 (--s). Lastly, for the R10.3 datasets, the medaka model was specified by using –model r103_min_high_g345, which was the only available for R10.3.

## Acknowledgements

We would like to thank John T. Longino and Michael Branstetter for providing valuable comments on the manuscript. For the Palaearctic phorid samples, we would like to thank Dave Karlsson, the Swedish Insect Inventory Project, and the crew at Station Linné that sorted out the phorids. We would also like to thank Wan Ting Lee for help with molecular work, and the numerous staff, students and interns who have contributed to the establishment of the pipeline in the NUS laboratory. We would also like to acknowledge Suphavilai Chayaporn and Niranjan Nagarajan from Genome Institute of Singapore for their help with basecalling using bonito. This work was supported by a Ministry of Education grant on biodiversity discovery (R-154-000-A22-112).

## Declarations: Software and test dataset availability

ONTbarcoder is available at https://github.com/asrivathsan/ONTbarcoder, which also contains the link to download the raw data and demultiplexing files. The manual for the software is included in the repository https://github.com/asrivathsan/ONTbarcoder/blob/main/ONTBarcoder_manual.pdf. The videos tutorials can be found in the YouTube channel Integrative Biodiversity Discovery: https://www.youtube.com/channel/UC1WowokomhQJRc71FmsUAcg. The datasets have been uploaded to NCBI, under BioProject: PRJNA745481, SRA accession numbers: SRR15185964, SRR15098600, SRR15098599, SRR15188571, SRR15188570,SRR15188569. The datasets, demultiplexing files and reference barcodes are also available via doi:10.5281/zenodo.5115258.

